# The impact of long-read sequencing on fungal genome assemblies: progress and disparity

**DOI:** 10.64898/2026.05.12.724544

**Authors:** E. Kroll, Y.A.B. Zoclanclounon, M. Urban, R. Hill, K.E. Hammond-Kosack

## Abstract

Fungal genomics has expanded rapidly over the past 30 years, and recently the pace and breath has further quickened for many taxa, although many taxonomic gaps persist. With three decades of rapid growth, fungal genomics now merits a re-examination of its history, progress, and unresolved taxonomic gaps. Here, we review the development of fungal genomics from early efforts such as the Fungal Genome Initiative to current progress driven by third-generation long-read sequencing. We have compiled and summarised publicly available fungal genomes to highlight trends in assembly quality, adoption of long-read technologies, and taxonomic representation. Notably, substantial phylogenetic gaps remain, particularly outside *Dikarya*, and significant challenges persist for unculturable taxa. This review identifies priorities for the fungal community, including: (1) coordinated efforts to close major taxonomic gaps across the fungal tree of life; (2) improved repository metrics to facilitate identification of high-quality assemblies; and (3) improved and standardised genome annotation which is lacking for most assemblies. Together, these steps will support the development of reliable genomic resources that capture the full breadth of diversity across the fungal kingdom, generating foundational data for comparative genomics, evolutionary biology, functional studies, genetic studies and applied research.

## Introduction

Over the past three decades, fungal genomics has expanded rapidly. The landmark publication of the *Saccharomyces cerevisiae* genome assembly in 1996^1^, was followed by the 2002 Fungal Genome Initiative (FGI) who proposed the sequencing of 15 fungal genomes carefully selected to represent key human and plant pathogens, model organisms, and evolutionary lineages^2^. Since then, the number of publicly available fungal genomes has expanded dramatically, with resources such as NCBI GenBank^3^ and the JGI MycoCosm portal^4^ now hosting >22,000 unique assemblies across the two databases. While assemblies now far exceed the 2002 targets, the field’s direction is now driven by the broader fungal community rather than a central body, creating a need for a resource to coordinate these distributed efforts.

The recent advent and maturation of high-throughput long-read sequencing technologies have resulted in dramatic improvements in genome quality across all domains of life. Despite comprehensive assessments of genome quality existing for plants^5^, animals^6^ and even insects^7^, no equivalent assessment of publicly available assemblies exists for fungi. This absence reflects the broader reality that fungi remain comparatively understudied relative to plants and animals, occupying a phylogenetically broad but historically neglected branch of the eukaryotic tree. Despite this fungi play central roles in global nutrient cycles, human and plant disease, industrial biotechnology, and microbiome function, and are critical to “One Health” frameworks^8,9^. The availability of high-quality, representative fungal genomes is foundational for comparative genomics, evolutionary biology, functional studies, genetic studies and applied research. Therefore, a consolidated understanding of how genome assembly data is distributed across the fungal tree of life, where major gaps remain, and how long-read sequencing has reshaped assembly completeness and accuracy is an essential resource to progress fungal research at a global scale.

In this review, we provide a high-level assessment of the current landscape of fungal genome sequencing, with a particular focus on the transformative impact of long-read sequencing technologies. Here we map the taxonomic distribution of available assemblies, evaluate their quality, and highlight persistent challenges and opportunities for the future of fungal genomics research.

### Landmark Genome Projects

Following the publication of the budding yeast *Saccharomyces cerevisiae* 12.1 Mb genome in (1996), there was a gap of six years before another model yeast and the first model filamentous genomes were published, namely the fission yeast *Schizosaccharomyces pombe* (2002) and the model filamentous fungus *Neurospora crassa* (2003), all using Sanger sequencing. In the same timeframe, 50-plus bacterial genomes were published. To rectify this serious imbalance the Fungal Genome Initiative emerged in 2002 led by Bruce Birren, Gerry Fink, and Eric Lander (https://www.genome.gov/Pages/Research/Sequencing/SeqProposals/FungalInitiative_Genome.pdf) to ensure that a global community strategy involving 15 carefully selected species (as described above) was adopted^2,10^. Of these, the genomes for nine species at pseudo-chromosome scale were published by 2007, another five by 2012 and the rest by 2015 (Fig. 1). By the end of 2012, before long-read sequencing became available, more than 100 reference fungal genomes from different species had been deposited in public repositories; many assemblies already included longer 454 sequencing reads ranging from ∼200 to 700 bp. In addition, many of these reference assemblies were already underpinning community resources by 2012. In 2016 and 2018, additional kingdom-wide efforts were instigated, namely the Kew Fungal Tree of Life Project (https://eetrust.org/projects/discovering-the-fungal-tree-of-life/) and the JGI 1000 Fungal Genomes Project (http://1000.fungalgenomes.org/) which aimed to fill the remaining phylogenetic gaps. However, progress towards achieving this aim has remained slow. In 2024, Kew’s Fungarium Sequencing Project commenced with the goal of sequencing > 2000 historical specimens, many being species types, from the world’s largest fungal collection.

**Fig 1.**
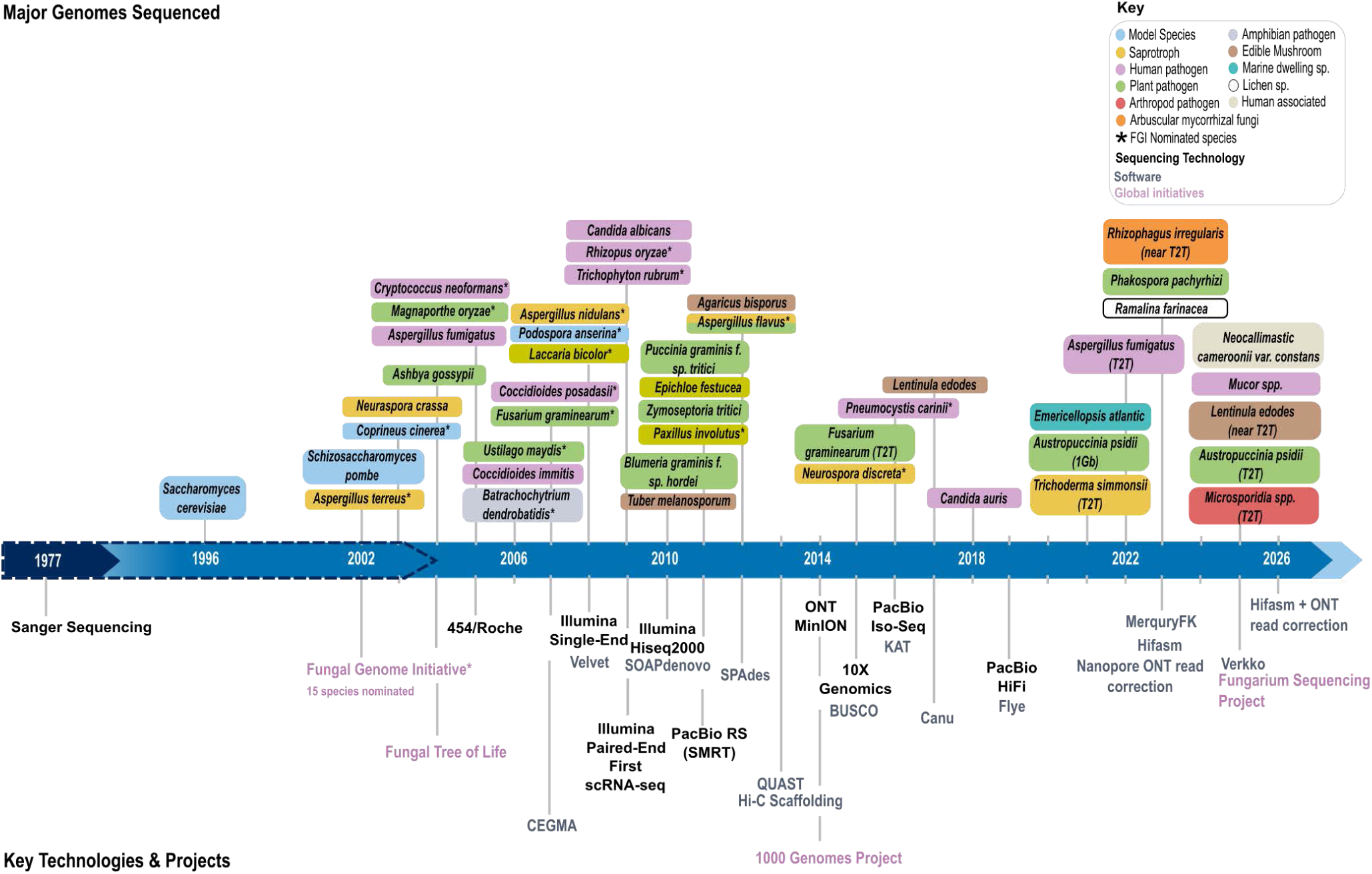
Timeline of key milestones in fungal genome sequencing. Timeline showing landmark fungal genome assemblies alongside the emergence of major sequencing technologies, community initiatives and analytical software. The dashed blue arrow highlights that Sanger sequencing was used to sequence all fungal genomes sequenced before 2004.

### Assembly Quality and Completeness

To evaluate the current landscape of fungal genomics, we curated a dataset of all publicly available genomes. Fungal genome assemblies were retrieved from NCBI^3^ and Joint Genome Institute MycoCosm^4^ databases on October 27, 2025, yielding a non-redundant and curated dataset of 22,799 genomes (Supplementary File 1, Fig. 2). Of these, 4,563 were generated using long-read sequencing technologies and 18,236 were generated using short-read sequencing platforms. Long-read assemblies demonstrated significantly higher contiguity, with a median contig N50 of 2.40 Mbp compared to 0.28 Mbp for short-read assemblies (p < 0.001; Fig. 2), with the upward trajectory in contig N50 from approximately 2020 onwards reflecting the progressive and widespread adoption of long-read sequencing technologies in fungal genomics (Fig. 3).

**Fig 2.**
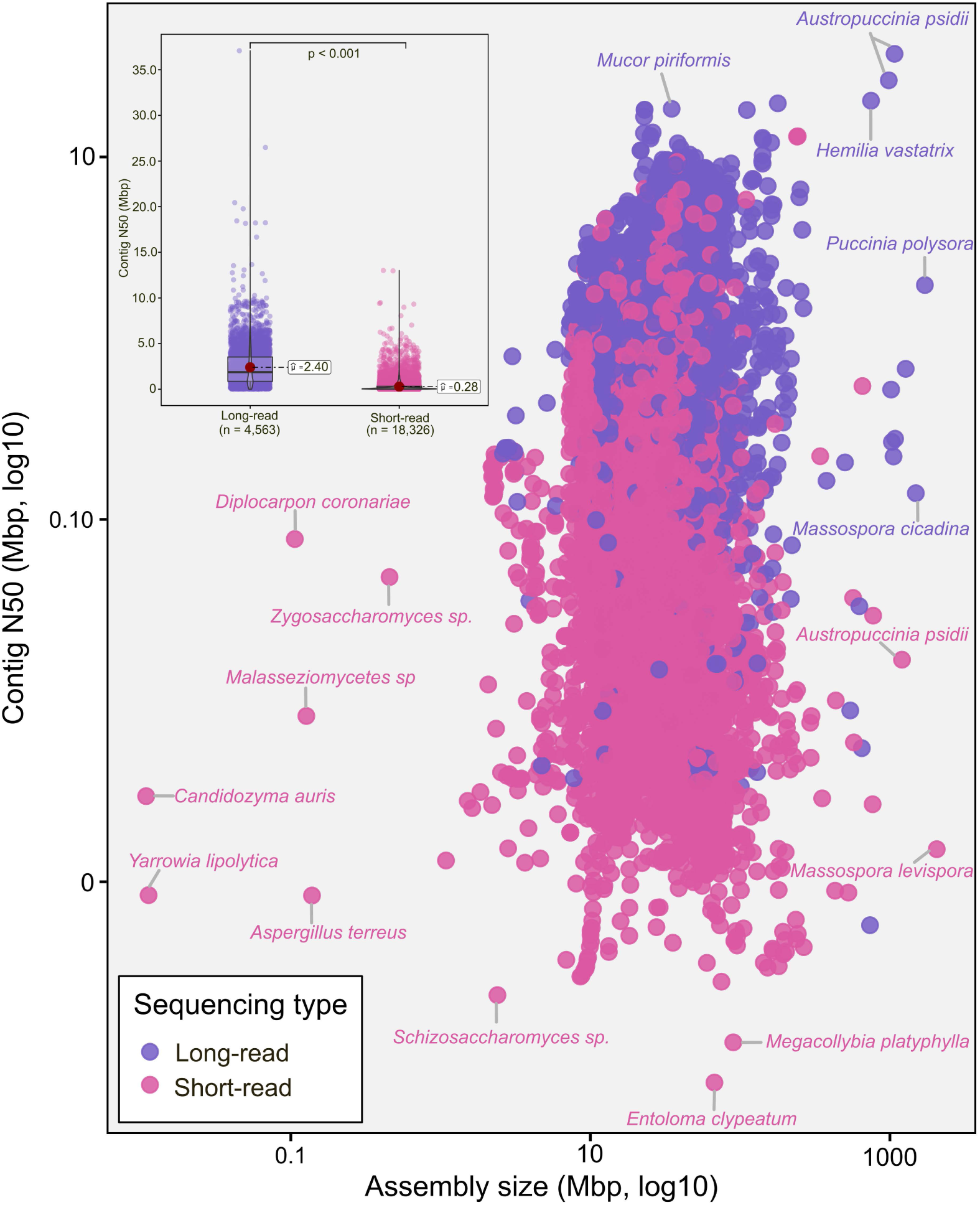
Relationship between assembly size and contig N50 across 22,799 fungal genome assemblies. Scatter plot of contig N50 (Mbp, log10) against total assembly size (Mbp, log10) for fungal genome assemblies generated from long-read (purple; *n* = 4,563) or short-read (pink; *n* = 18,326) sequencing platforms. Inset: box plots comparing contig N50 distributions between sequencing categories; horizontal dashed lines indicate median values (long-read: 2.40 Mbp; short-read: 0.28 Mbp), and filled circles denote means. Statistical differences between groups were assessed using Welch’s one-way ANOVA followed by Games-Howell post-hoc test.

**Fig 3.**
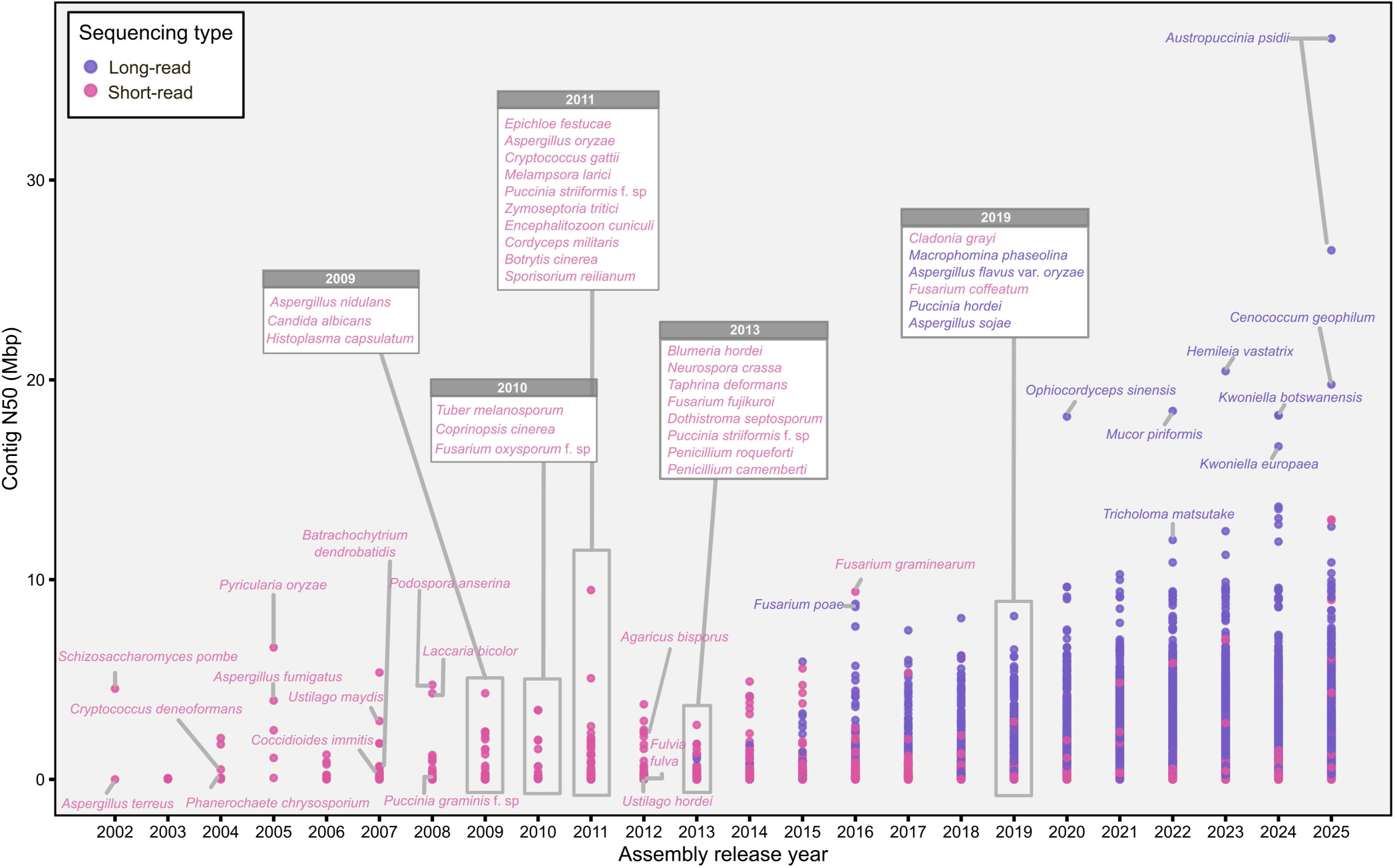
Trends in assembly contiguity across fungal genome assemblies released between 2002 and 2025. Contig N50 values (Mbp) plotted against assembly release year for long-read (purple) and short-read (pink) fungal genome assemblies. Boxed annotations and labelled assemblies represent historically significant or contiguity-notable genomes within each release year. The first Saccharomyces genome sequenced in 1996 and 42 low-quality assemblies (released between 1997 and 2001) flagged by NCBI (Supplementary File 1) were excluded from this plot.

Genome completeness was assessed using BUSCO^11^ and varied across both sequencing categories (Fig. 4). Notably, assemblies of *Hanseniaspora* spp. and microsporidian species formed distinct low-completeness clusters (Fig. 4a), attributable to their compacted or evolutionarily divergent genomes^12–15^. Among assemblies with contig N50 < 0.1 Mbp, *Candida* spp., *Saccharomyces* spp., *Hanseniaspora* spp., and *Nematocida* spp. exhibited a distinctive pattern of similar BUSCO completeness values within this low-contiguity fraction (Fig. 4b).

**Fig 4.**
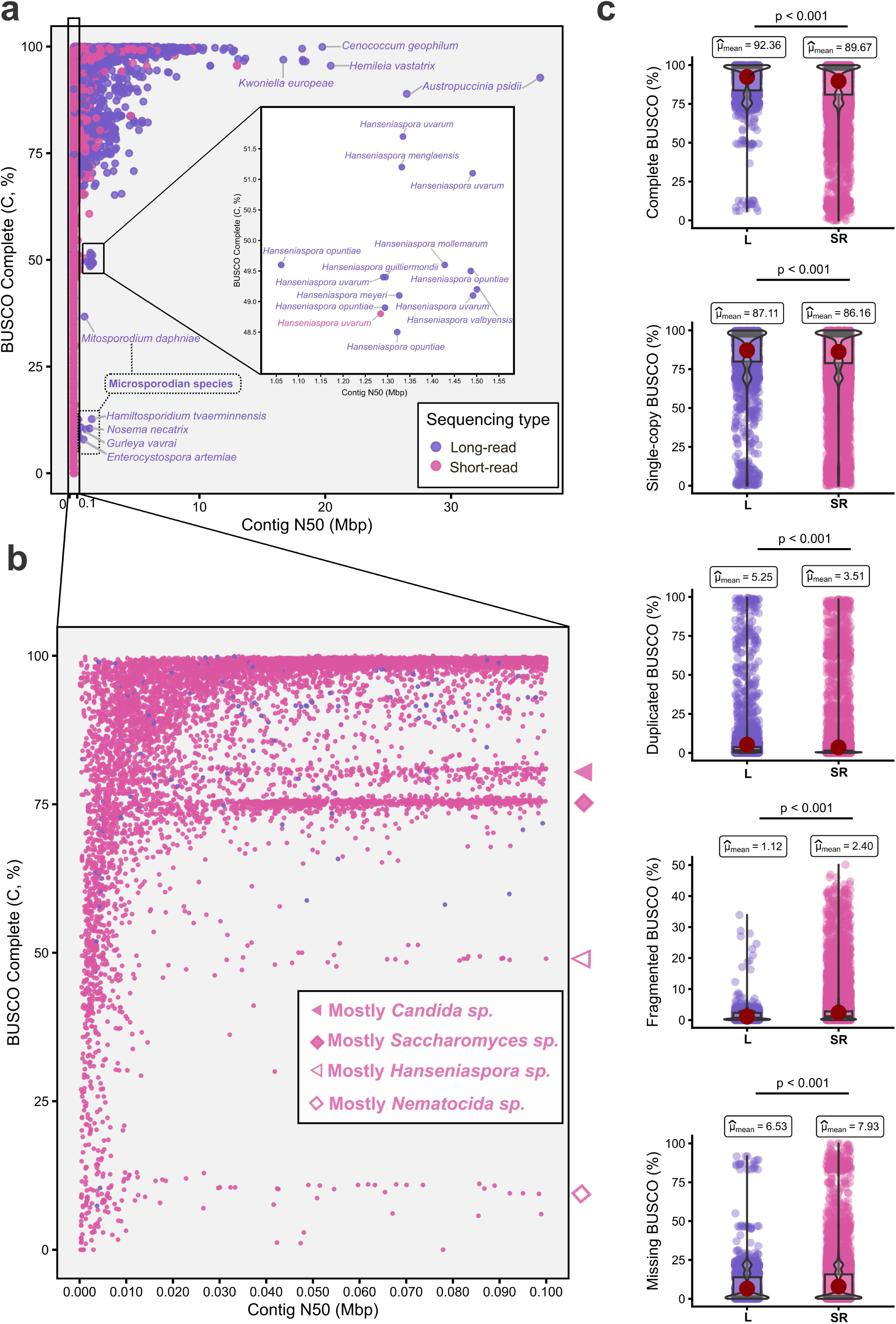
BUSCO completeness metrics across 22,799 fungal genome assemblies stratified by sequencing technology. (a) Complete BUSCO scores (%) plotted against contig N50 (Mbp) for all assemblies. Inset enlargements highlight clusters of Hanseniaspora spp. (upper inset) and microsporidian species (lower inset), which exhibit characteristically reduced BUSCO completeness attributable to their highly compacted or evolutionarily divergent genomes. (b) Expanded view of assemblies with contig N50 < 0.1 Mbp; horizontal markers denote species-level clusters dominated by *Candida* spp., *Saccharomyces* spp., *Hanseniaspora* spp., and *Nematocida* spp., which display similar BUSCO completeness values within this low-contiguity fraction. (c) Violin and box plots comparing distributions of complete, single-copy, duplicated, fragmented, and missing BUSCO percentages between long-read (L) and short-read (SR) assemblies. All pairwise comparisons were statistically significant (p < 0.001, Games-Howell post-hoc test). Sequencing type is indicated by colour throughout (long-read: purple; short-read: pink).

As expected, long-read assemblies achieved a higher mean complete BUSCO score of 92.36% versus 89.67% for short-read assemblies (p < 0.001; Fig. 4c) yet exhibited elevated duplication rates (mean 5.25% versus 3.51%), potentially reflecting unresolved haplotypes arising from residual heterozygosity or true gene expansion^16,17^. Conversely, short-read assemblies exhibited higher fragmentation (2.40% versus 1.12%) and missing BUSCO proportions (7.93% versus 6.53%), likely reflecting the limited capacity of short reads to resolve repetitive and structurally complex genomic regions. Overall, long-read sequencing confers measurable improvements in both contiguity and gene-space recovery across fungal genomes.

### Taxonomic Coverage and Gaps

Most assembled fungal genomes fall in the subkingdom *Dikarya*, especially lineages of yeasts (*Saccharomycetes*), plant pathogens (*Sordariomycetes*) and ubiquitous air and soilborne microfungi (*Eurotiomycetes*) in the phylum *Ascomycota*; and commonly observed macrofungi (*Agaricomycetes*) and opportunistic human pathogen yeasts (*Tremellomycetes*) in the *Basidiomycota* (Fig. 5a). Other basidiomycete lineages known for smut (*Ustilaginomycetes*) and rust (*Pucciniomycetes*) plant pathogens are similarly more highly represented.

**Fig 5.**
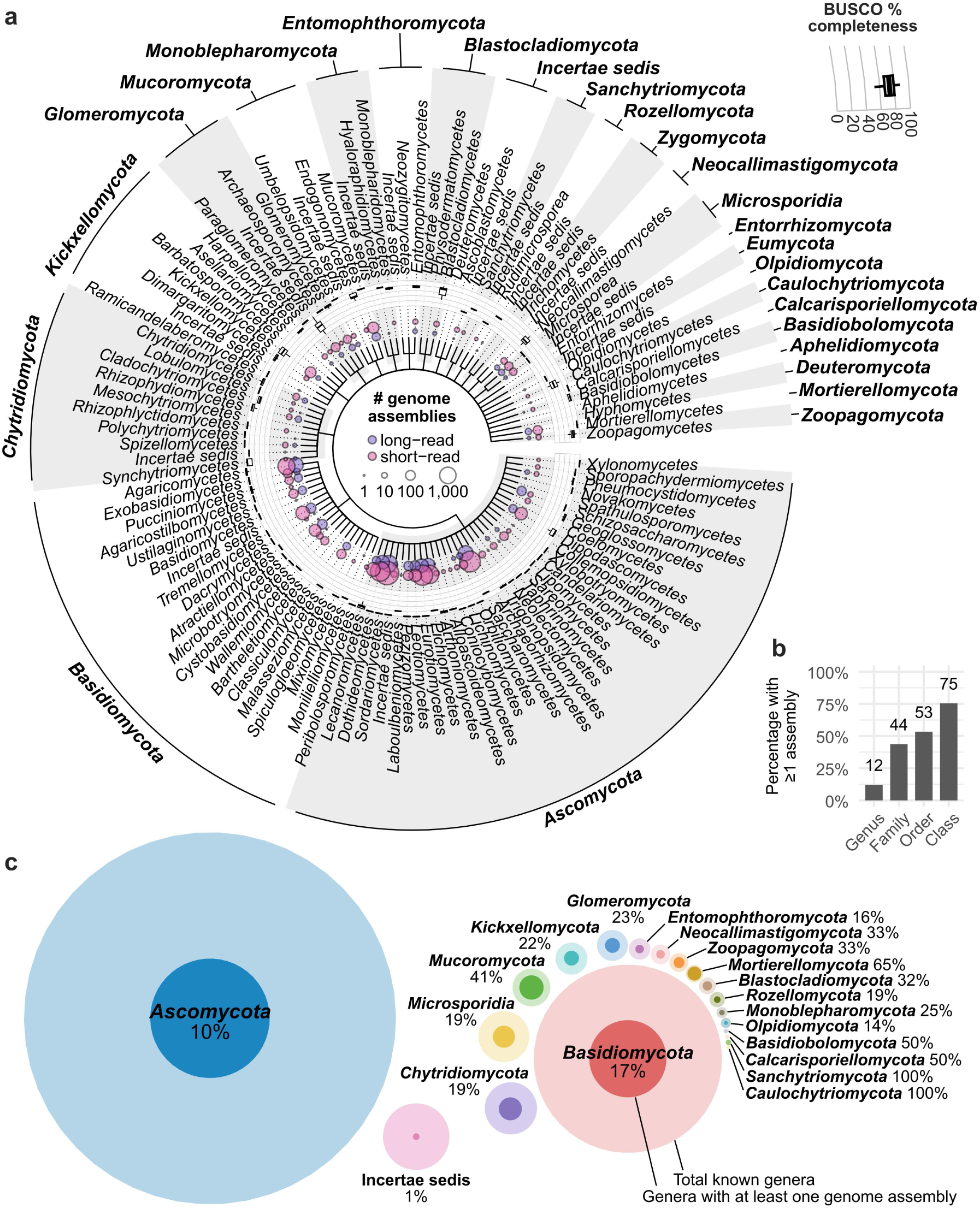
Taxonomic distribution of fungal genome assemblies deposited in NCBI and JGI MycoCosm. Taxonomy is assigned according to MycoBank (as of 20/10/2025). **(a)** Taxonomic tree with tree tips representing the rank of class. For a detailed order-level tree see Supplementary Fig. 1. The inner ring shows the total number of short- and long-read assemblies and the outer ring shows the percentage BUSCO completeness per class. **(b)** The percentage of taxa at each taxonomic rank with at least one representative genome assembly. **(c)** The percentage of genera within each phylum with at least one representative genome assembly.

Biases in sequencing efforts towards economically or clinically important fungi, or those that are easier to sequence, are to be expected, but they limit our deeper understanding of genomic diversity across the kingdom. Even within the better studied *Dikarya*, there are still notable gaps. The *Sordariomycetes* and *Dothideomycetes* are two of the largest classes in the *Ascomycota* and are among the best represented in terms of total number of genome assemblies, and yet many orders still have no representative assembly (Fig. 5a); overall 53% of fungal orders and only 12% of fungal genera have a representative genome assembly (Fig. 5b). In fact, only 10% of ascomycete and 17% of basidiomycete genera have at least one genome assembly (Fig. 5c), emphasising the major work that remains. Initiatives to generate genomic resources across the fungal tree of life are crucial to address these gaps (e.g. the ongoing 1000 Fungal Genomes Project, Varga et al. 2025^18^, The Darwin Tree of Life Project Consortium 2022^19^).

Predictably, taxonomic coverage of long-read genome assemblies mirrors that of short-read, although in some cases the number of long-read assemblies has caught up or even exceeds the number of short-read assemblies (e.g. *Schizosaccharomycetes, Ascomycota; Microbotryales, Microbotryomycetes, Basidiomycota*), demonstrating the rapid uptake of these technologies. Although non-dikaryotic phyla such as *Glomeromycota* and *Mucoromycota* have previously lagged in terms of genomic resources, growing representation for these lineages is now evident, although challenges remain especially for uncultivable, unicellular non-*Dikarya* such as *Rozellomycota*^20^. A key factor impeding the transition to long-read WGS in such lineages is the difficulty in obtaining sufficient DNA, as the conventional practice of culturing the fungus to obtain high molecular weight (HMW) DNA is not possible. This instead requires whole genome amplification, which comes with its own biases and complexities^21^.

### DNA extraction methods and emerging technologies for recalcitrant fungi

High purity DNA is essential for long-read sequencing, as contaminants inhibit enzymatic library preparation and reduce sequencing efficiency. Long-read sequencing approaches that enable near telomere-to-telomere (T2T) assemblies broadly depend on preparation of high-molecular-weight (HMW) DNA, which is in practise typically obtained through culturing. Platforms such as PacBio HiFi and Oxford Nanopore typically require input fragments in the ∼48-50 kb range, where read length and DNA integrity directly determine assembly contiguity and resolution of repetitive regions. Sequencing providers therefore impose strict quality thresholds on DNA samples, including fragment size distributions, prescribed UV light absorbance ratios and fluorometric quantification to monitor DNA sample quality.

In fungal DNA extractions, frequently obtained low A260/230 ratios indicate co-purified contaminants that lead to reduced read length, lower yield, and compromised assembly quality^22^. As a result, sequencing centres often waive data output guarantees. A wide variation in fungal genome size exists with 2-3 Mb in Microsporidia to large >1 GB repeat-rich genomes in obligate biotrophic rust fungi requiring further adaptation of protocols^23^. Dikaryotic and multikaryotic states in some taxa complicate haplotype resolution, increasing the requirement for high-quality input DNA and tailored assembly strategies^24,25^. Recent studies show that recovery of HMW DNA is highly sensitive to growth stage, biomass, and extraction chemistry while harvesting mycelium prior to accumulation of melanin and other secondary metabolites can markedly improve DNA quality and sequencing performance^26^. Key trade-offs are that low-input DNA extraction methods using small DNA amounts suit slow-growing strains but often reduce purity, whereas approaches using more biomass produce longer fragments. These differences directly influence fragment length distributions and, ultimately, assembly contiguity.

Classical extraction methods, including CTAB and phenol-chloroform protocols, remain widely used, often with modifications to minimise DNA shearing and remove inhibitors^22^. These protocols are often cost-effective and adaptable across species, although optimisation is typically required. Commercial kits (e.g. Cytiva Nucleon PhytoPure Genomic DNA Extraction Kit, Macherey-Nagel NucleoBond HMW Kit) have also been adapted successfully for fungal DNA extraction, with variable performance depending on species and input material^26^. Downstream size-selection tools such as BluePippin (Sage Science) are commonly used to enrich long fragments for long-read sequencing. No single protocol is universally effective, and success remains species dependent.

A major limitation in fungal genomics is the large proportion of unculturable taxa, including obligate biotrophs (e.g. powdery mildews, rusts) with low biomass production and slow-growing symbionts such as lichen-forming fungi^27^. Culture-independent approaches are essential for these organisms. Emerging methods like single-cell sequencing are increasingly bypassing the need for traditional cultivation^28^.

Metagenomic sequencing of host-associated material enables bioinformatic genome reconstruction directly from environmental samples, although contamination and binning accuracy remain limiting factors. The lichen-forming fungus *Solorina crocea* was assembled from metagenomic data, demonstrating the feasibility of genome recovery from obligate symbionts^29^. Single-cell DNA sequencing approaches based on microfluidics or fluorescence-activated cell sorting (FACS) of nuclei can reduce sample complexity and enables sequencing from low-input or mixed samples in non-dikaryotic parasitic fungi^30^. The various described methods are beginning to unlock previously inaccessible organism diversity.

### Genome Structure

Genome assembly is undergoing a major advancement from highly fragmented draft approximations toward chromosome-complete and increasingly telomere-to-telomere (T2T) representations of entire genomes. Advances in long-read sequencing, particularly highly accurate single-molecule reads and ultra-long sequencing, have made it feasible to resolve repetitive regions that historically fragmented assemblies, including centromeres, subtelomeres, segmental duplications, and large structural variants. As a result, completeness is no longer defined primarily using coarse proxies such as BUSCO, but by the presence of full-length gapless chromosomes including centromeres and bounded by telomeric sequence^31^. This transition reframes reference genomes from approximations of gene space to comprehensive maps of genome architecture, enabling more accurate comparative genomics, structural variation analysis, and evolutionary inference across diverse taxa, and establishing T2T assemblies as an emerging gold standard^31,32^.

One aspect of fungal genome organisation that has been unlocked by long-read sequencing is the improved detection of accessory chromosomes. These are chromosomes that are non-essential but may carry advantageous genes, and can vary in presence across individuals^33^. Although typically smaller than core chromosomes, as accessory chromosomes are generally highly repetitive, long-read sequencing has greatly improved their recovery during assembly. While accessory chromosomes have previously been associated mostly with pathogens, not all pathogens have accessory chromosomes (e.g. Cuomo et al., 2007^34^; Hill et al. 2025^35^) and not all fungi with accessory chromosomes are pathogens (e.g. Zhang et al. 2026^36^). Nonetheless, accessory chromosomes have been most frequently observed in pathogens in the *Ascomycota*, which may partially be explained by the sequencing bias for this group (Fig. 3), as robust delimitation of accessory chromosomes requires chromosome-level assemblies of multiple individuals of a species. Examples of accessory chromosomes carrying key pathogenicity factors were initially established for specific *Fusarium* and *Alternaria* fungi, namely *Fusarium oxysporum* f. sp. *lycopersic*i where 25% of the genome is accessory and many proteinaceous effector SIX genes reside on accessory chromosomes^37^, and *Alternaria alternata* f. sp. *mali* where accessory chromosomes carry toxin biosynthesis genes such as AM-toxin genes conferring pathogenicity on apple trees^38^. The loss of accessory chromosomes can even induce a pathogenic-to-mutualistic lifestyle switch, as shown in the poplar pathogen *Stagonosporopsis rhizophilae*^39^.

Accessory chromosomes are one manifestation of the broader concept of genome compartmentalisation, where different genomic regions exhibit different architectures, either as distinct accessory chromosomes or within core chromosomes. Genome compartmentalisation has typically been defined from regions showing greater colocalisation of repetitive content and effector proteins that are involved in host-fungal interactions^40^. The successful resolution of these repetitive regions has again been advanced by highly contiguous assemblies thanks to long-read sequencing. Such repeat- and effector-rich compartments have been observed in certain plant pathogens^41^, animal pathogens^42^ and mycorrhizal fungi^43^. How and why specific fungi have evolved such compartments while others in similar ecological niches have not is still not understood, but repeat-rich regions are broadly understood to be origins for adaptive genome evolution^44^.

Transposable elements (TEs) have long been understood to influence genome architecture thanks to their ability to move and independently replicate within the genome^45^. Long-read sequencing has revolutionised TE biology in fungi by enabling the discovery of a new superfamily of giant TEs, *Starships*, specific to the species-rich subphylum *Pezizomycotina*^46^. These *Starships* are referred to as ‘cargo-mobilising’, as they include up to hundreds of miscellaneous accessory genes which can seemingly move around the host genome or even horizontally between individuals or species^47^.

Similar to accessory chromosomes, *Starship* identification, gain and loss is contingent on long-read sequencing of multiple individuals of the same species, to not only span their length (up to 700 kbp^48^) but also detect large insertions relative to other genomes. As a result, *Starships* have predominantly been identified from well-studied lineages such as *Aspergillus*, *Penicillium* and *Fusarium*^49^ but more widespread long-read sequencing will doubtless unveil the true extent and impacts of these giant TEs.

### Fungal pangenomes

The first formalised concept of the pangenome^50^, refers to the full complement of genes across all individual species, including both the conserved core genome and the variable accessory genome. Across fungi, pangenome resources are now available for a rapidly growing but still taxonomically uneven set of species, dominated by *Ascomycota* and a smaller number of *Basidiomycota*, with genome counts ranging from a handful of isolates to well over a thousand in intensively studied taxa including industrial (*Saccharomyces)* and human-associated taxa (*Candida*, *Aspergillus*) and a broad range of plant pathogens spanning major cereal, legume, fruit, and tree-infecting genera (e.g., *Zymoseptoria, Fusarium, Rhizoctonia, Pyrenophora, Parastagonospora, Claviceps, Magnaporthe,* and *Colletotrichum*) (Supplementary File 2). Most fungal pangenomes, built from short-read drafts, rely on reference-anchored comparisons that underestimate structural variation and accessory chromosomes.

Pangenome graphs provide a unified genomic framework that represents sequence diversity across individuals while reducing the reference bias inherent to single linear genomes. By encoding alternative alleles, presence–absence variation, and large structural variants as parallel paths, graph-based pangenomes enable more accurate read mapping and variant discovery, particularly in repetitive or highly divergent genomic regions^51,52^. Graph representations further allow scalable integration of new genomes and support comparative and association studies across diverse lineages using a shared coordinate system. Within current fungal pangenome studies, graph-based pangenome resources remain rare but are now emerging in both filamentous ascomycetes and budding yeast, and these advances are tightly coupled to the availability of near-complete, long-read–enabled assemblies. In the *Fusarium oxysporum* species complex, graph pangenomes have been reported alongside long-read assemblies that better resolve accessory chromosomes and lineage-specific regions that are typically collapsed or fragmented in short-read drafts^53,54^. In the cereal pathogen *Pyrenophora tritici-repentis*, a graph-based pangenome was similarly built from (partial) long-read assemblies to capture structural diversity beyond SNP-scale variation^55^. In yeasts, graph-based representations have been constructed for *Saccharomyces cerevisiae* using long-read assemblies^56^, and at larger scale in a recent study using 1086 near T2T assemblies^57^.

Across these examples, T2T assemblies, or at a minimum long-read assemblies approaching T2T contiguity, are effectively prerequisites for high-fidelity pangenome graph construction in fungi to avoid spurious graph branching driven by assembly fragmentation, collapsed repeats, or mis-localised paralogs, and to place structural-variant breakpoints accurately in coordinate systems shared across isolates^58^.

### Improved Transcriptomics and Gene Models

While long-read sequencing has been most widely adopted for fungal genome assembly, it is increasingly transforming fungal genome annotation and functional genomics through long-read RNA sequencing (lrRNA-seq). By capturing full-length transcripts, lrRNA-seq overcomes isoform-reconstruction ambiguities inherent to short-read RNA-seq, which are exacerbated in fungi by compact genes, short introns, overlapping transcription, and condition-dependent transcript boundaries^59,60^. Two experimental strategies dominate: PacBio Iso-Seq, which provides high-accuracy full-length cDNA sequences, and Oxford Nanopore Technologies (ONT) cDNA or native (direct) RNA sequencing, which offers flexible throughput and, in the case of direct RNA, access to native RNA features.

To date, the most widespread application of lrRNA-seq in non-yeast fungi has been improved genome annotation. Integrating Iso-Seq with short-read RNA-seq has led to substantial revisions of gene models, including corrected exon–intron structures, extended untranslated regions (UTRs), alternative transcription start and termination sites, and extensive alternative splicing. This has been demonstrated across diverse filamentous fungi, including the wheat pathogens *Zymoseptoria tritici*^60^, and *Fusarium graminearum*^61^, the medicinal mushroom *Ganoderma lingzhi*^62^, the filamentous ascomycete *Aspergillus flavus*^63^, the entomopathogen *Cordyceps militaris*^64^, the rust fungus *Phakopsora pachyrhizi*^65^, and the model basidiomycete *Coprinopsis cinerea*^66^. Collectively, these studies reveal far greater transcriptomic complexity than predicted *ab initio*, including pervasive alternative splicing, widespread read-through transcription and polycistronic transcripts, and thousands of previously unannotated genes. While these studies highlight the potential of lrRNA-seq, the complexity of lrRNA-seq is accompanied by a suite of challenges. These include its higher error rate relative to short-read sequencing, the presence of isofrom artefacts, and the immaturity of the current annotation tools to handle lrRNA-seq input^67^.

### Conclusions and Future Perspectives

Fungal genomics began over 30 years ago with a small number of foundational reference genomes, but the field has since expanded toward capturing the extraordinary diversity of the fungal kingdom. Yet despite this rapid growth, progress remains constrained by a landscape in which high-quality assemblies sit alongside fragmented genomes, annotations are often sparse or inconsistent, and sequencing efforts remain disproportionately focused on a narrow set of tractable or economically important taxa. These limitations hinder our ability to study fungal biology and fungal genetics, as low quality genomes and annotations can conceal lineage-specific innovations, and could impede our ability to detect the genomic evolution underpinning ecological transitions^68,69^, host specialisation^70,71^, and pathogenic emergence^72^.

The most pressing challenge is the phylogenetic imbalance in publicly available genomes. The overwhelming majority are derived from the *Dikarya*, concentrated in *Saccharomycetes*, *Sordariomycetes*, *Eurotiomycetes*, and a subset of Basidiomycota such as *Agaricomycetes* and *Tremellomycetes*. Groups with major ecological, human health, industrial, or agricultural relevance, including the difficult dikaryotic rusts and smuts, are comparatively well represented, but large groups of the fungal tree of life remain genomically invisible (Fig 3). This bias reflects a strong emphasis on economically or clinically important fungi, as well as those that are more tractable for sequencing, but this persisting bias substantially limits our understanding of genomic diversity across the kingdom. Targeted efforts to sample across the full phylogenetic breadth of the fungal kingdom do exist, i.e. the Kew’s Fungal Tree of Life Project started in 2015, and the JGI’s 1000 Fungal Genomes Project started in 2018, but without greater community efforts and uptake, our understanding of fungal evolution will remain fundamentally incomplete.

Equally urgent is the need for consistent, high quality genome annotation. In total, only a quarter of fungal genome assemblies in NCBI are annotated to date. Although the number of assemblies is rapidly increasing year on year, the proportion of those assemblies with annotated gene models is, in fact, decreasing (Supplementary Fig. 2), showing that genome annotation efforts are not keeping pace with sequencing and assembly. As discussed above, lrRNA-seq can provide a source of evidence to improve the quality and completeness of genome annotation, however annotation workflows remain a bottleneck due to both computational and biological complexity. In addition to transcriptomic evidence, gold-standard annotation also uses known proteins/gene models from related species and probabilistic models to integrate these diverse sources of evidence into gene predictions (e.g. Gabriel et al. 2024^73^). New deep learning tools trained on existing gene models show promise for rapid annotation^74,75^, however their performance relative to conventional workflows has not been comprehensively benchmarked for fungi, especially lesser studied non-dikaryotic lineages. Furthermore, current repositories lack metrics on the availability of annotations, annotation methods, completeness, and quality warnings. In addition, different annotation methods can substantially distort comparative analyses, from orthogroup inference to pangenome construction. Evidence from a pan-annotation study in *Fusarium*^76^, underscores how variation in annotation can skew and misdirect comparative conclusions between genomes. Establishing community standards for annotation workflows, metadata reporting, and independent quality assessment must therefore become a priority if fungal genomics is to enable robust and reproducible insights.

Even with improved annotation, a deeper conceptual challenge remains, the very definition of the genome in a complex organism. In many biological systems, the genome is implicitly treated as a single, stable sequence associated with an individual organism. In fungi, however, this assumption is frequently violated by the dynamics and plasticity of fungal genomes. Chromosome copy number variation, including pervasive and often reversible aneuploidy, can arise rapidly and be stably maintained over ecologically or clinically relevant timescales, blurring the boundary between genotype and karyotypic state^77^. Heterokaryotic species further complicate this picture as they harbour multiple genetically distinct nuclei within a shared cytoplasm. These complex genomic states then intersect with the extensive structural variation provided by highly variable and lineage-specific accessory chromosomes^78^. Together, these features constitute major drivers of fungal genome plasticity and underpin their capacity to adapt rapidly to environmental and selective pressures^79^.

Resolving this complexity requires a shift towards phased, chromosome-level assemblies which are essential for fully resolving multinucleate and heterokaryotic assemblies. This would allow us to better understand how known plasticity-generating mechanisms, such as transposable-element mobilisation, region-specific mutation rates, and the gain or loss of accessory chromosomes, operate within the true biological context of multinucleate fungi, offering a far more accurate understanding of the evolutionary and economic consequences of fungal genomic diversity. Yet current genome repositories rarely capture this complexity. Databases typically provide a single, collapsed assembly per isolate and lack standardised metadata on chromosome counts, structural compartmentalisation, or heterokaryotic status. Incorporating such metadata through explicit reporting of nuclear composition, ploidy variation, and accessory chromosome complements would markedly improve the interpretability of fungal genomes.

Although long-read sequencing has revolutionised fungal genome assembly, many fundamental aspects of fungal genome biology operate at scales beyond the reach of single-molecule reads. This includes long-range scaffolding technologies (e.g. Hi-C/Omni-C, Strand-seq, and optical mapping)^80,81^, which provide genome-wide linkage and spatial interaction information that can be used to order, orient, and phase contigs into chromosome-scale assemblies, thereby resolving large-scale structural organisation and repeat-rich regions that remain inaccessible to long reads alone. A further advancement is the emerging field of 3D genomic architecture^82^. 3D genome studies in fungi reveal diverse higher-order organization, including bipartite compartmentalisation^83,84^, self-interacting domains^83,84^, centromere clustering^85^, and the spatial segregation of accessory chromosomes^84^ across multiple fungal lineages.

Taken together, the field is moving toward a future in which fungal genomics is defined not by isolated reference genomes, but by comprehensive, phylogenetically balanced, population-scale resources. Fungal genomes expanded from 20 to over 20,000 in the past thirty years, shifting the primary challenge from data generation to interpretation.

In this Review, we have synthesised current metrics of genome quality across publicly available fungal genomes and, in doing so, have found deficiencies in transparency, standardisation, and annotation availability. Addressing these gaps will be essential for transforming the growing volume of genomic data into robust, biologically meaningful insight. Ultimately, the next phase of fungal genomics will depend not only on expanding dataset size, but on improving data quality, accessibility, and interpretability to enable high-quality science.

## Acknowledgements

K.H.K., M.U., E.K, and R.H. were supported by the Biotechnology and Biological Sciences Research Council (BBSRC) Institute Strategic Programme (ISP) grant Delivering Sustainable Wheat (BB/X011003/1) within the work package Delivering Resilience to Biotic Stress (BBS/E/ER/230003B Earlham Institute and BBS/E/RH/230001B Rothamsted Research). Y.A.B.Z. was supported by the BBSRC grant Green Engineering (BB/X010988/1).

## Contributions

All authors contributed to conceptualisation, writing, development and revision of this manuscript. E.K. and Y.A.B.Z. curated and analysed the data. R.H. mapped the data to taxonomic frameworks and contributed substantially to discussion of content. K.H.K. and M.U. provided key contributions to the historical context of fungal genomics.

## Supplementary information

**Supplementary Fig. 1.** Detailed order-level taxonomic distribution of fungal genome assemblies deposited in NCBI and JGI MycoCosm.

**Supplementary Fig. 2.** The proportion of fungal genome assemblies in NCBI that have annotations available.

**Supplementary File 1. Genome assemblies and BUSCO quality assessments.** Lists all fungal genome assemblies included in this study. Genomes were obtained from NCBI and JGI as of 27 October 2025. Only published and unrestricted genomes from JGI were included in the analysis to respect JGI’s data use policies regarding unpublished projects. For each assembly, metadata including accession number, assembly name, organism and strain, sequencing technology, and assembly statistics are reported. Genome completeness and quality were assessed uniformly using BUSCO fungiodb (n = 1,122), with counts of complete (single-copy and duplicated), fragmented, and missing orthologs provided for each genome.

**Supplementary File 2. Published fungal pangenome studies.** A summary of published fungal pangenome studies, reporting the analysed species or species complex, higher-level clade, number of genomes included, and the corresponding reference. The file also indicates whether long-read assemblies and graph-based pangenome approaches were used.

## Data Availability

All metadata associated with this study are provided in Supplementary File 1, which also includes accession numbers for all genome assemblies. Genome assemblies and their associated publications can be accessed via NCBI Datasets (https://www.ncbi.nlm.nih.gov/datasets/genome/) and JGI MycoCosm (https://mycocosm.jgi.doe.gov/mycocosm/home).

## Code Availability

All code used in this study has been deposited in a public GitHub repository at https://github.com/Yedomon/fungal_genome_assemblies, with a permanent archived copy available on Zenodo (https://zenodo.org/records/19923620).

