## Supplementary figures and images for "The impact of long-read sequencing on fungal genome assemblies: progress and disparity"

### Supplementary Fig. 1

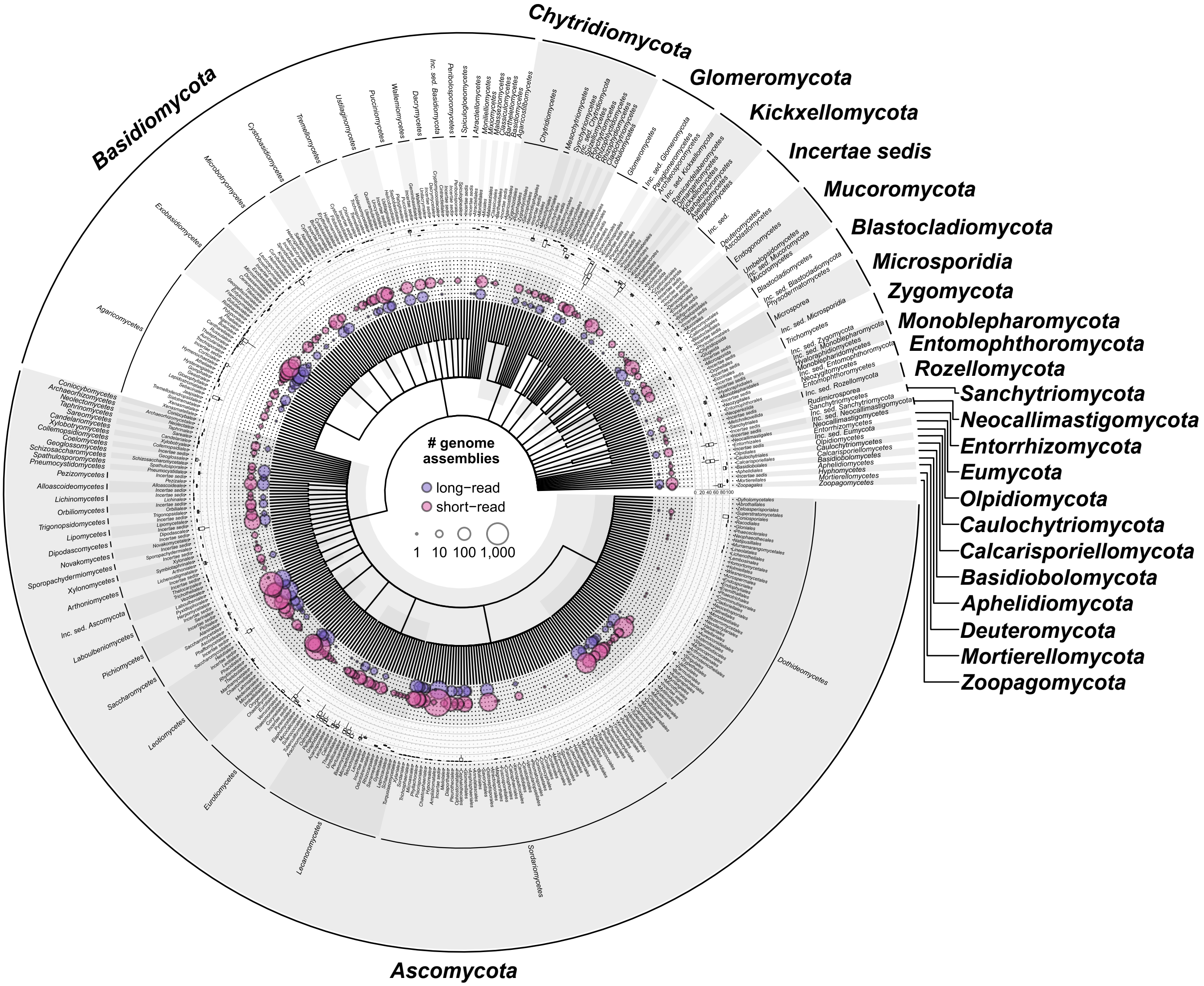

### Supplementary Fig. 2

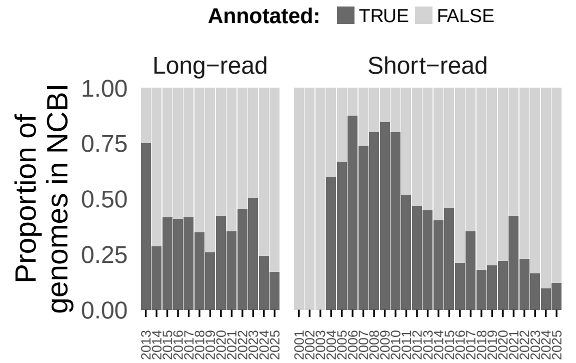
